# Island biogeography through the lens of multiscale metapopulation dynamics: insights into species-area relationships

**DOI:** 10.1101/2025.03.21.644586

**Authors:** Maxime Clenet, François Munoz, Coline Picard, Adam Bernard, Estelle Pitard

## Abstract

While island biogeography focuses on species richness equilibrium driven by immigration and extinction, and metapopulation theory examines single-species dynamics across fragmented habitats, their interplay remains poorly understood. In particular, the species-area relationship remains a subject of ongoing debate, yet there are limited theoretical foundations to explain it. To address this, we developed a multiscale stochastic metapopulation model to investigate diversity patterns on islands, bridging the gap between island biogeography and metapopulation theory. Our model integrates regional colonization from a mainland with local colonization-extinction processes within islands at the single-species level, then extends this to multiple, independent species. By analyzing the stationary properties of this model, we generate novel predictions of Species Area Relationship (SAR) based on local extinction rates, within-island colonization rates, and mainland immigration rates. We demonstrate how the interplay of these parameters influences the relationship, predicting patterns that can resemble either the power-law of Arrhenius or the semi-logarithmic relationship of Gleason, depending on the relative importance of mainland immigration versus within-island dynamics, and on the nature of the species abundance distribution in the mainland. This unified framework offers new insights into the mechanisms driving species richness and distribution across spatial scales, providing a more holistic understanding of biodiversity patterns in fragmented landscapes.

## 1 Introduction

The Species-Area Relationship (SAR) is a cornerstone pattern of major influential theories both in ecology and biogeography. Many phenomenological models have been proposed to describe SAR, and have been compared and challenged using different datasets and statistical approaches [41]. The most classical, prominent “law” was proposed first by Arrhenius [2] in the form of a power law,

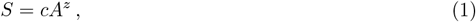

where *S* represents the number of species, *A* the island area, *c* a constant, and *z* the exponent that characterizes the particular group of organisms under consideration. The exponent *z* is typically less than 1 [8], indicating that the increase in species richness slows down as the area expands. Since then, this relationship has become a foundational concept in ecology, though it has been subject to refinement and debates [23]. The main concurrent view, proposed by Gleason [13], suggests that the SAR should be described by a semi-logarithmic model rather than a power law. According to Gleason’s law, the SAR is expressed as:

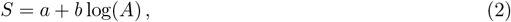

where *a* and *b* are constants. Here, *a* represents the baseline species richness, while *b* reflects the sensitivity of species richness to changes in area. Both parameters depend on factors such as regional species pool, habitat type, and the kind of organism considered. This model predicts that species richness increases logarithmically with area, meaning that smaller areas experience a more pronounced increase in species richness with changes in area, while larger areas show a slower rate of increase in richness as the area expands further. Gleason’s law challenges the traditional power-law relationship of Arrhenius law, suggesting a more gradual and less steep increase in species richness as area increases. The debate between these two formulations—logarithmic versus power-law scaling— has persisted in biogeography, with different datasets providing support for one or the other [5, 37]. Despite the rich empirical studies exploring these patterns and some variation in the theoretical formula of SAR [41], there remains a notable gap in theoretical models of species dynamics that could predict such biogeographical “laws”. Most research to date has concentrated on describing observed relationships rather than developing mechanistic frameworks to explain the underlying processes. Bridging this gap is crucial for advancing our understanding of how species richness responds to changes in habitat area. A deeper theoretical foundation could not only shed light on the drivers of these patterns but also help resolve the debate between competing models, providing a unified framework for addressing species-area dynamics.

There are many contexts in ecology where we want to compare richness or other biodiversity patterns for varying sampling area. Here we specifically consider the context of island biogeography theory [26, 27], in which islands are distinct and isolated pieces of land that can exchange migrants between them and/or with a continent. Island biogeography has received broad interest in surveys of biodiversity not only on oceanic islands, but also over a broad spectrum of “habitat islands”, i.e., fragments of suitable habitat surrounded by a matrix of unsuitable environmental conditions [27, 43]. With such scope, island biogeography has inspired theory and practices in biological conservation [43], such as the well-known SLOSS (Single Large Or Several Small) framework [7], which guides the design of reserve networks. We refer to Island Species Area Relationship (ISAR) to characterize species richness-area relationship among islands. In their foundational theory, McArthur and Wilson [27] posited that species richness in islands results from a dynamical balance between immigration from outside the islands and extinction events inside. The most classical case is the mainland-island model, such as each island can receive immigrants from a common mainland source. Immigration rates can decrease with the distance from the source continent, while extinction rate should increase in smaller islands. The fundamental idea is that when richness increases in an island, the number of species that can go extinct increases proportionally, while less new immigrant species can establish. This yields a “dynamical” equilibrium state such as the number of extinction and immigration events are balanced and an equilibrium species richness is reached in the island. This is called “dynamical” because, while richness is expected to be stationary, the identity of the species varies as some go extinct, while other immigrate. The two major ecological mechanisms are thereby the immigration and the extinction processes. Because these processes depend on island size and isolation from the mainland, the equilibrium richness varies accordingly and can give rise to the famous ISAR *S* = *cA*^*z*^. Here we propose to extend the framework to acknowledge the multiscale nature of immigration and extinction dynamics in metapopulations, while considering that different species individually and independently undergo metapopulation dynamics.

Metapopulation theory offers a relevant framework by emphasizing how colonization and extinction dynamics shape individual species distribution across fragmented habitats [22, 15]. This framework accounts for variation in population density and occurrence within a network of habitat fragments, providing insights into the interplay between local and regional processes. Because of the shared focus on immigration and extinction dynamics, the connection between island biogeography and metapopulation dynamics has been addressed, although quite rarely [34, 16]. For instance, Ovaskainen & Hanski [34] used the Incidence Function metapopulation model to characterize the local probability of presence depending on the connectivity and the occupancy of potential sources. As in the model we will present, the authors assumed independent dynamics of different species in a common habitat. In their case, the island includes a single local population, and island area influences both colonization and extinction probabilities. Then they considered that the potential richness of a habitat patch is the sum of the probability of presence for each species. However, when considering a large array of island sizes in biogeography, it may be unrealistic to consider that a vast island includes a single population of a species. Therefore, we propose a multiscale metapopulation extension in which an island contains an internal metapopulation governed by colonization and extinction dynamics. External colonization can also occur from the other metapopulation islands [20], or from a mainland source [1].

In this paper, we introduce the multiscale metapopulation model to explore diversity patterns in islands. By integrating island biogeography and metapopulation dynamics, we provide novel insights into the mechanisms driving ISAR. In line with the island biogeography theory, we designed a “mainland-island” model, which assumes an infinite mainland where species persist indefinitely and occasionally send colonizers to islands within which finite metapopulation dynamics occur [20, 1]. We define the island size as the number of habitat patches in the island, and island richness as the number of species that can maintain at least one population in the island metapopulation. We expose analytical and simulation-based properties of this model, and propose a simple multispecies generalization by considering that several species undergo metapopulation dynamics.

We provide novel predictions of ISAR in islands based on three fundamental parameters: the local population extinction rate, the within-island colonization rate and the mainland-island colonization rate. We posit that these parameters determine an equilibrium probability distribution for the number of populations in the island, allowing us to infer species richness within the island. We discuss in which conditions the predicted ISAR can better fit one or another phenomenological relationship. Specifically, we show that ISAR can be very similar to either the Arrhenius or the Gleason law depending on the model parameter values.

## 2 Material and methods

### 2.1 Multiscale mainland-island metapopulation model

The seminal metapopulation model, first proposed by Levins [22], describes the colonization and extinction dynamics of species within a patchy environment. Building on this model, subsequent extensions incorporated immigration from a persistent mainland source, thereby giving rise to the mainland-island metapopulation model [18, 15] (see App. B and Fig. 1a). These deterministic models assume an infinite number of habitat patches, which, while analytically convenient, limits their applicability to real-world systems composed of a finite number of patches. A key consequence of this assumption is the exclusion of stochastic extinctions, which occur in finite systems due to random fluctuations in metapopulation dynamics [9]. Even when the colonization and extinction rates favor species persistence (*c > e*), finite populations remain vulnerable to extinction due to demographic stochasticity. The importance of stochasticity in island biogeography was already recognized by MacArthur and Wilson [26], who proposed a stochastic mainland-island model to describe the balance between immigration, local extinction, and species persistence on islands.

**Figure 1:**
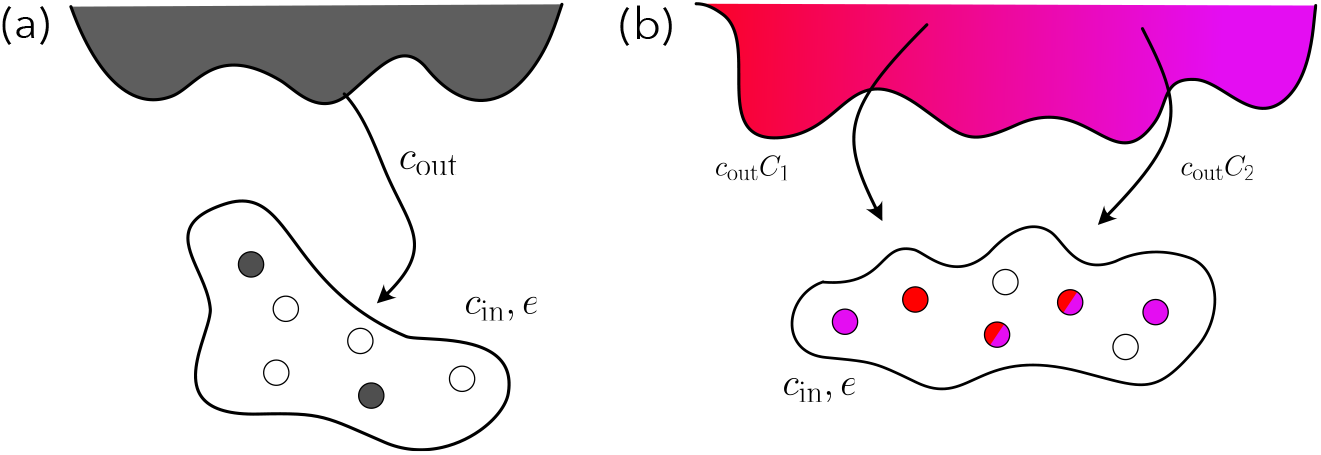
Scheme of the mainland-island model for one species (a) and two species (b). In Fig. (a), the scheme illustrates the dynamics of a single species present on the mainland. The colonization rate from the mainland to the island is denoted by *c*_out_. The island is divided into a set of habitat patches, which can be either occupied (filled circles) or empty (void circles). The local dynamics on the island are governed by the colonization rate *c*_in_ and the extinction rate *e*. In Fig. (b), the model is extended to two species. The colonization rate from the mainland becomes *c*_out_*C*_*m*_, where *C*_*m*_ depends on the relative abundance of the species on the mainland. On the island, both species follow the same dynamics and can coexist in the same habitat patches.

In this paper, we present a multiscale extension of the stochastic mainland-island model, integrating colonization-extinction dynamics across two spatial scales: a local scale, where species occupy a network of habitat patches on the island, and a regional scale, where species disperse from the mainland to the island. To explore these dynamics, we consider a simplified framework with a single island (see Fig. 1). Specifically, we analyze a multispecies version of the model, where an island consists of *n* habitat patches and supports a set of *M* species. Each species, indexed by *m* ∈ 1, …, *M*, is treated as independent, following its own colonization-extinction dynamics within the island.

A common approach for modeling stochastic metapopulation dynamics is through a continuous-time Markov chain. The state of the system is defined by the number of occupied patches *k*, with transitions between states occurring randomly over time. The transition rates between states of species *m* depend on two key processes represented by 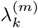 and 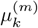, for all *k* ∈ [0, *n*]. First, the rate at which new species occupy empty patches on the island is given by

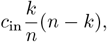

where *c*_in_ is the within-island colonization rate, allowing colonization from occupied patches to empty patches. Second, the immigration rate from the mainland to the island is given by

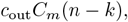

where *c*_out_ is the mainland-island colonization rate, which can reflect the distance between the mainland and the island. For each species *m*, the number of occupied patches, *k*, changes according to the following state transitions:

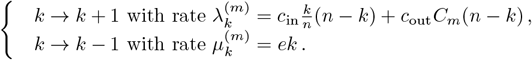

The parameter *e* represents the extinction rate of an occupied patch on the island and is assumed to be independent of the island size *n*.

Each species immigrates from the mainland at the same baseline rate *c*_out_, but its actual colonization rate depends on its relative abundance in the mainland. We denote the relative abundance of species *m* in the mainland as *a*_*m*_, and its corresponding relative abundance as *C*_*m*_, where:

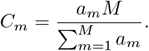

This formulation ensures that species with higher abundance in the mainland have a proportionally higher probability of colonizing the island (see Fig. 1b for an illustration with two species).

The distribution of species abundances in the mainland, *P* (*a*_*m*_), is generally unknown. A common simplifying assumption is that all species have equal relative abundance, meaning that each species has the same likelihood of colonizing the island. This assumption can be modeled using a discrete uniform distribution, where the abundance of any species *X* in the mainland follows:

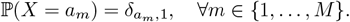

Under this uniform scenario, each species has an equal abundance *a*_*m*_ = 1, leading to *C*_*m*_ = 1, meaning all species have the same mainland-island colonization rate.

Alternatively, species abundances may follow a heterogeneous distribution, such as the widely used logarithmic series distribution (LSD), introduced by Fisher *et al*. [10]. The LSD is commonly applied in ecology to describe species relative abundances. It is defined by a shape parameter *θ* ∈ (0, 1), and the probability mass function is:

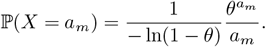

In this case, species with higher abundance in the mainland have greater colonization rates on the island, contributing more significantly to the colonization process, while rarer species are less likely to reach and establish.

The dynamics of species *m* is governed by a set of master equations, i.e., a system of differential equations that describes the probability 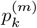 that *k* patches are occupied by species *m*:

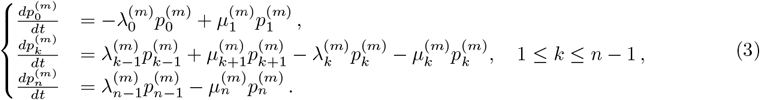

### 2.2 Equilibrium properties

Alonso and McKane [1] derived the existence and uniqueness of the equilibrium distribution of model

(3) in the single-species case (see A.C). Let *p*_*k*_ denote the probability that *k* patches are occupied. The equilibrium of model (3), denoted as *p*_*k*_(∞), can be derived analytically by setting the time derivative to zero 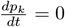. Assuming *c*_out_ ≠0, it has been demonstrated [1] that the equilibrium distribution is given by:

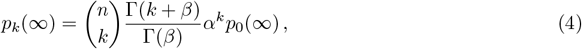

where Γ(*x*) is the gamma function, 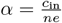 is a dimensionless parameter representing the strength of the colonization rate within the island relative to the local extinction rate, and 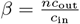 is a dimensionless parameter representing the input of immigrants from the mainland relative to colonization within the island.

Given that the sum of probabilities over all states is 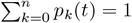,the probability of having zero occupied patches at equilibrium, *p*_0_(∞), is given by:

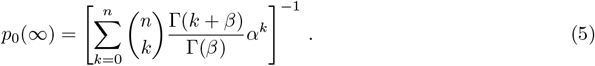

Figure 2 shows the equilibrium distribution *p*_*k*_(∞) as a function of *k*, obtained through numerical integration of Eq. (4). For sufficiently large *c*_out_, the distribution approximates a Gaussian shape, a result that can be formally derived using the Van Kampen expansion [21], a method for approximating solutions to master equations in large populations. For small *c*_out_ (hence small *β*), the probability of extinction *p*_0_(∞) increases proportionally to 1*/β* (see A.D), indicating a high risk of metapopulation collapse.

**Figure 2:**
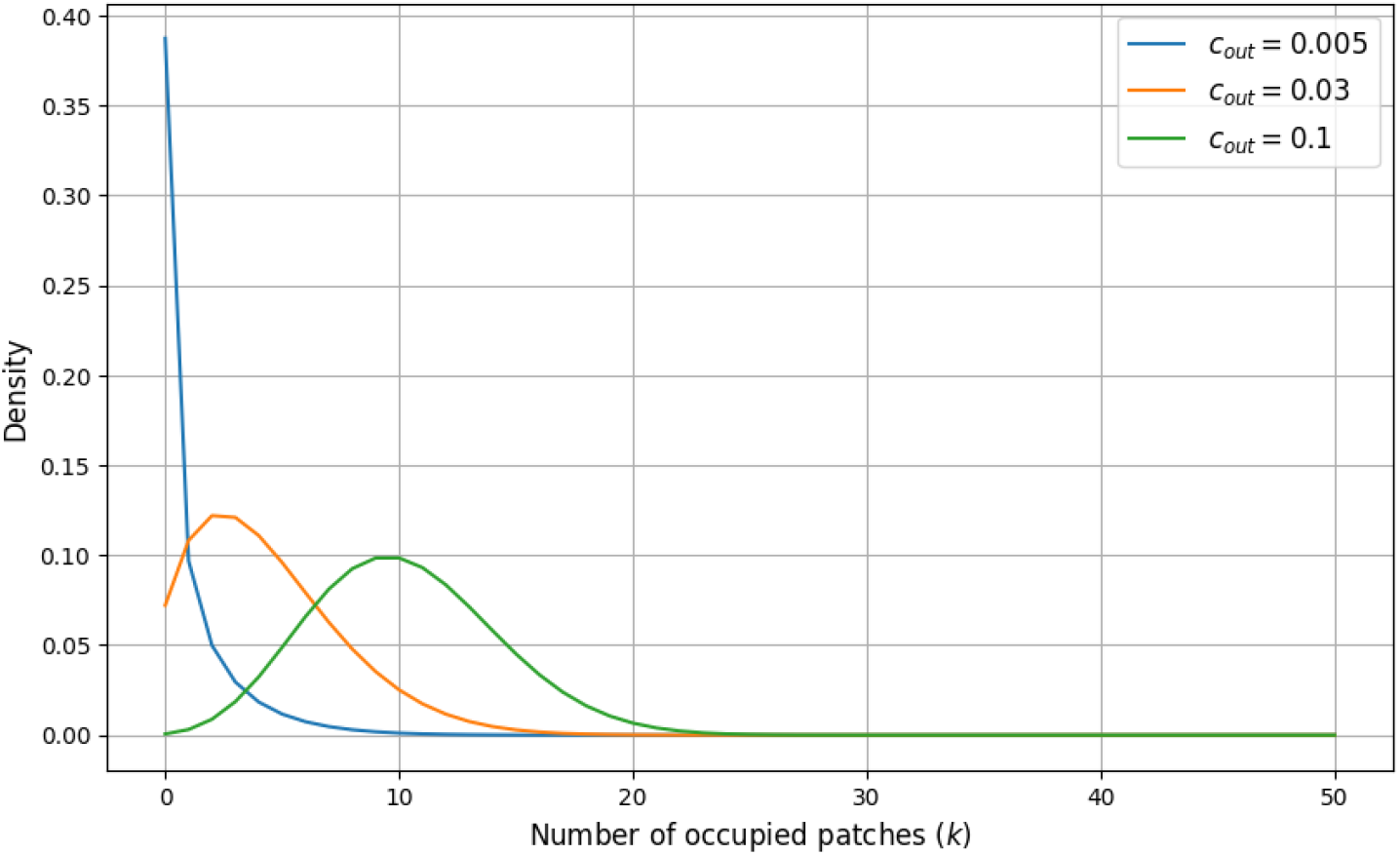
Equilibrium distribution *p*_*k*_(∞) for different mainland-island colonization rates (*c*_out_) with a within-island colonization rate *c*_in_ = 0.8, a local extinction rate *e* = 1, and an island size *n* = 50. The blue line represents the distribution for *c*_out_ = 0.005, showing a sharp peak at *k* = 0, indicating a high probability of having zero occupied patches. The orange line, with *c*_out_ = 0.03, shows a more spread-out distribution, peaking at a higher number of occupied patches. The green line, for *c*_out_ = 0.1, depicts a broader distribution that approaches a Gaussian-like shape, peaking at an even higher number of occupied patches. These distributions highlight how increasing the colonization rate from the mainland shifts the equilibrium towards higher occupancy levels in islands, reflecting the balance between colonization and extinction processes.

Empirical studies on plant dispersal [31] suggest that long-distance colonization events are rare, supporting the assumption that *c*_out_ is generally much smaller than the within-island colonization rate *c*_in_ and that island sizes *n* remain limited, ensuring *β* stays small. In the extreme case where *c*_out_ = 0, the model simplifies to a logistic stochastic system, exhibiting distinct dynamics such as a quasi-stationary state (see A.D for further details on these limiting cases).

### 2.3 Species richness and approximation of the probability of extinction

To study the ISAR, the key quantity of interest is species richness (*S*), which depends on the island area, here represented by the number of habitat patches *n*. Our goal is to determine its value in the equilibrium state as *t*→ ∞. Understanding these dynamics is crucial for predicting how species richness responds to ecological factors such as colonization and extinction rates.

In the equilibrium state, as *t* → ∞, species richness can be expressed as:

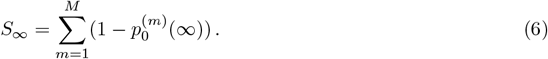

where 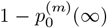 represents the probability that species *m* is present on the island at equilibrium (i.e., has at least one population). It provides a quantitative measure of long-term species richness on the island by leveraging the probability of extinction for each species.

This approach is advantageous because it incorporates both stochastic and deterministic aspects of the metapopulation model. It is directly linked to the species-level dynamics defined earlier, making it consistent with the underlying model assumptions. Additionally, this measure is robust across scenarios with different values of the parameters *e, c*_in_, and *c*_out_, providing a flexible tool for evaluating species richness under varying ecological conditions.

At large times (*t* → ∞), the asymptotic behavior of *p*_0_(*t*) for a single species depends on the relationship between the within-island colonization rate *c*_in_ and the extinction rate *e*. Recall that we consider the case *c*_out_ ≠ 0 yet *c*_out_ small. If *c*_in_ *< e*, under the approximation that the growth rate *λ*_*k*_ varies linearly with *k*, the master equation can be solved analytically using the method of characteristics (see A.E), leading to the following simplification:

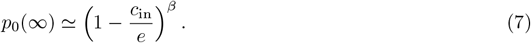

This expression smoothly approaches 1 as *c*_out_ → 0, indicating a high likelihood of the system remaining in state *k* = 0. If *c*_in_ ≥ *e*, the linear approximation is no longer valid. Since within-island colonization exceeds extinction and is supplemented by constant mainland immigration, we expect *p*_0_(∞) to approach zero (see [1] for a detailed derivation).

In the scenario where the species abundances on the mainland are heterogeneous, Eq. (6) can be generalized by replacing *c*_out_ with *c*_out_*C*_*m*_ in 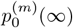,the asymptotic probability of extinction of species *m*, leading to the following approximation for *c*_in_ *< e*:

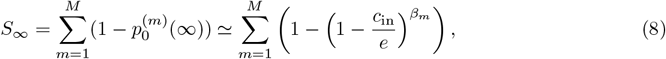

where 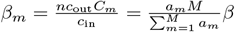. The approximation follows from Eq. (7).

## 3 Results

### 3.1 Homogeneous species abundance distribution in the mainland

Motivated by the biogeographic laws introduced earlier in the paper, we analyze the Island Species-Area Relationship (ISAR) using the multiscale metapopulation model (3). In this section, we consider the scenario where species are homogeneously distributed on the mainland, meaning that all species can arrive on the island at the same rate *c*_out_.

We focus on the regime where local extinction is possible, specifically when *c*_in_ *< e*. For our analysis, we set *c*_in_ = 0.8 and *e* = 1, ensuring that species populations go extinct locally in the absence of immigration from the mainland. To explore the behavior of the ISAR, we compute equilibrium species richness, *S*_∞_, as a function of island size *n* for different values of *c*_out_ (see Fig. 3a). When *c*_in_ *< e* and *β*≪ 1, species richness can be approximated using Eq. (8), which explicitly expresses *S*_∞_ as a function of *n*. This allows us to generate data points for varying island sizes and distances from the mainland. Our results indicate that ISAR exhibits different behaviors depending on *c*_out_, with species richness tending to saturate as island size increases (see Fig. 3a).

**Figure 3:**
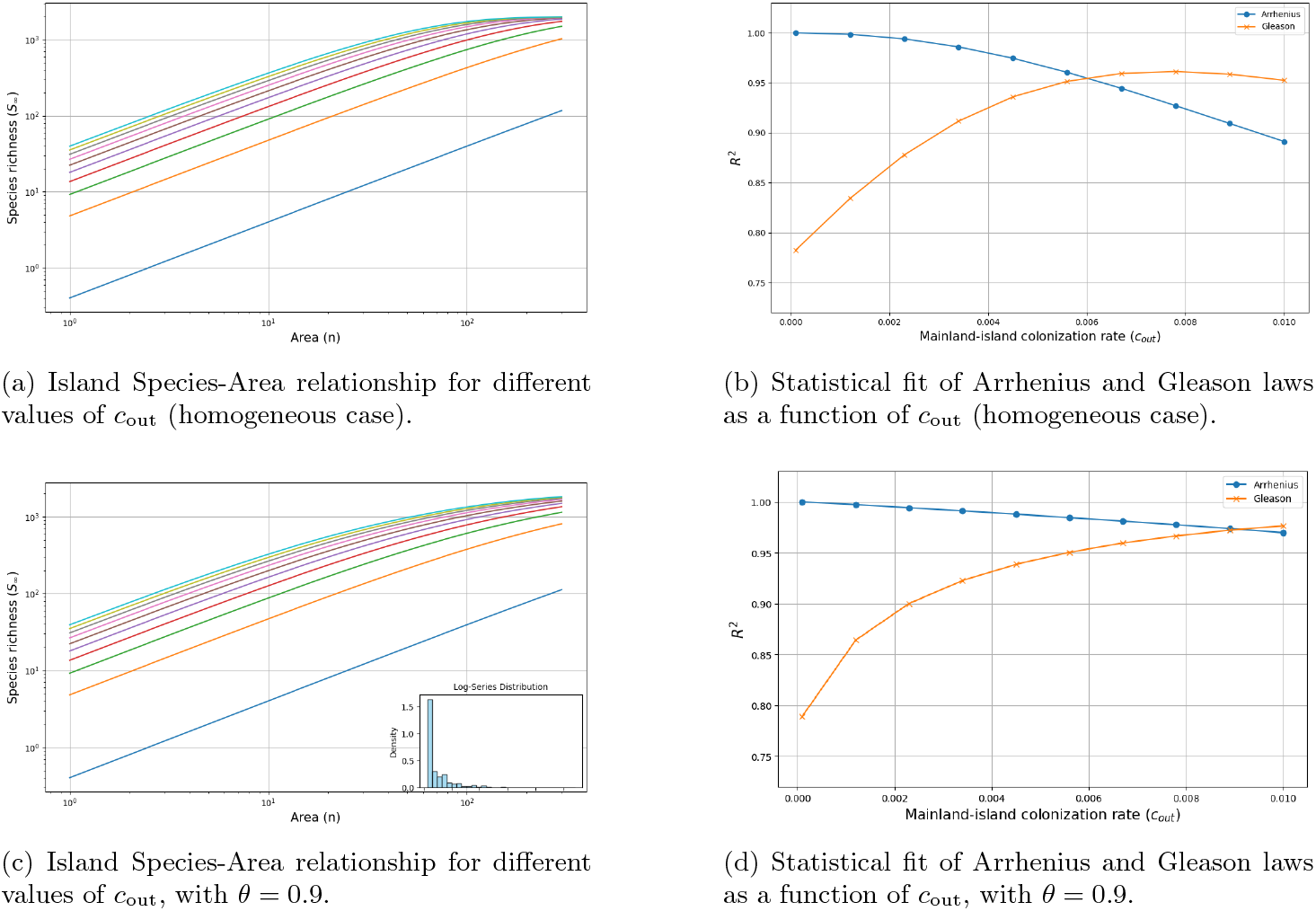
Prediction of species-area relationships with the multiscale metapopulation model (3) under homogeneous (a, b) and heterogeneous (c, d) species abundance distributions. In Fig. (a), Species-area relationship for different mainland-island colonization rates (*c*_out_), illustrating how species richness (*S*_∞_) varies with island area (*n*) when species abundances (*a*_*m*_) are uniform in the mainland. The parameters are *M* = 2000, *c*_in_ = 0.8, *e* = 1. The mainland-island colonization rate *c*_out_ varies from 10^−4^ (bottom curve) to 10^−2^ (top curve), highlighting how increasing dispersal from the mainland influences species accumulation on islands. In Fig. (b), the coefficient of determination *R*^2^ is shown for the statistical fit of the Arrhenius (1) (blue) and Gleason (2) (orange) laws to the predictions in Fig. (a), as a function of *c*_out_. Each point represents a specific mainland-island colonization rate, ranging from 10^−4^ to 10^−2^, illustrating how well these classical laws approximate the species-area patterns predicted by the multiscale metapopulation model in the homogeneous case. In Fig. (c), the same analysis as in (a) is performed but with a heterogeneous species abundance distribution in the mainland, following a Fisher log-series with parameter *θ* = 0.9. A subplot within (c) displays this distribution. The comparison between Figs. (a) and (c) highlights the effect of species abundance heterogeneity on the species-area relationship. In Fig. (d), the coefficient of determination (*R*^2^) is shown for the statistical fit of the Arrhenius and Gleason models to the predictions in (c), allowing direct comparison with (b) to assess the impact of species abundance heterogeneity on model performance.

Next, we compare the ISAR generated by our metapopulation model with two commonly used models: the Arrhenius power-law model (1) and the Gleason logarithmic model (2). Using a non-linear least squares fitting procedure (detailed in A.A), we quantify how well each model describes species richness patterns. As shown in Fig. 3b, the Arrhenius model provides an excellent fit for small values of *c*_out_. However, as island size increases, species richness eventually saturates due to the decreasing number of extinct species, reducing the agreement with a power-law relationship. Above a certain threshold of *c*_out_, the Gleason model provides a better fit than the Arrhenius model.

To further assess model accuracy, we examine confidence intervals for the exponent *z* in the Arrhenius model (see Fig. 4). As *c*_out_ increases, the exponent *z* decreases, and its confidence interval widens, indicating greater variability in species richness predictions. This aligns with the observed decreasing goodness-of-fit of the Arrhenius model as island size increases. Notably, the exponent *z* remains within the expected range, approximately between 0 and 1, reinforcing the idea that species richness on islands scales sub-linearly with area, but with increasing deviations as colonization rates grow.

**Figure 4:**
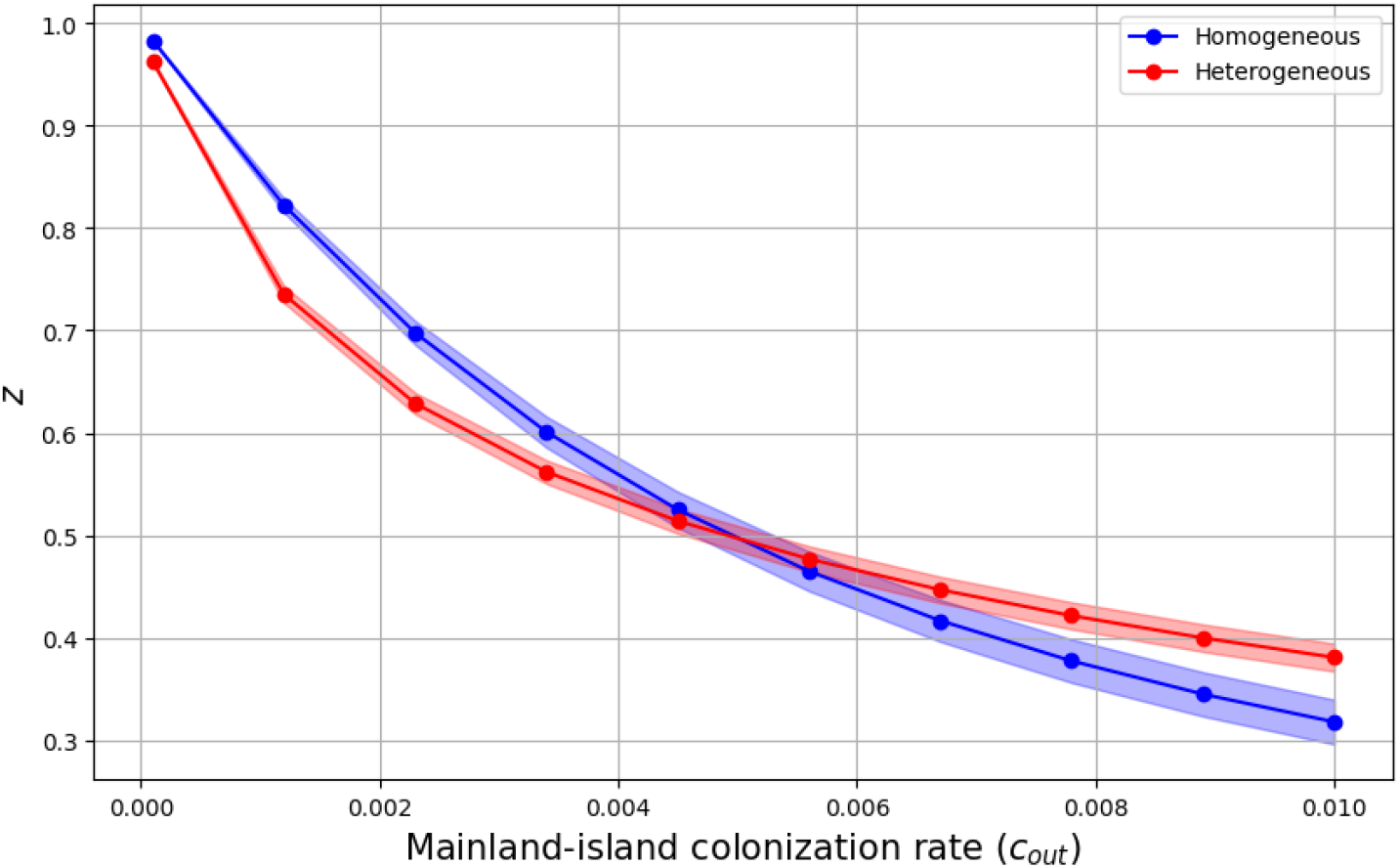
Arrhenius exponent *z* as a function of the mainland-island colonization rate *c*_out_. The exponent *z* is obtained from the statistical fit of the Arrhenius species-area relationship to the predictions shown in Figures 3a (homogeneous case) and 3c (heterogeneous case). The blue line represents the case of homogeneous species abundance, while the red line corresponds to the heterogeneous case, where species abundances follow a Fisher log-series distribution with *θ* = 0.9 in the mainland. For each estimated value of *z*, a 99% confidence interval is plotted, representing the statistical uncertainty in the exponent estimation. This interval quantifies the variability in the fit due to species abundance distributions and *c*_out_.

### 3.2 Heterogeneous species abundance distribution in mainland

When species abundances on the mainland follow a heterogeneous distribution, the resulting ISAR exhibits notable differences compared to the homogeneous case. To highlight this contrast, we consider an extreme scenario where species abundances are drawn from a logarithmic series distribution with *θ* = 0.9 (see Fig. 3c, subplot for an example of the distribution). As in the homogeneous case, we assume that island populations depend on immigration, such that *c*_in_ *< e*.

Using the same methodology as in the homogeneous case, we observe in Fig. 3c that the ISAR retains a similar shape across all values of *c*_out_, again displaying a saturating effect. However, in the heterogeneous case, saturation occurs at slightly larger island sizes, suggesting that species richness accumulates more gradually when mainland species abundances are unevenly distributed.

This effect is further highlighted in Fig. 3d, where the Arrhenius model maintains a strong fit over a broader range of island sizes before the Gleason model becomes the better descriptor. Additionally, compared to the homogeneous case, both the Arrhenius and Gleason models exhibit a better overall fit across all investigated values of *c*_out_. This suggests that heterogeneous species abundances contribute to a more predictable species-area relationship, possibly due to the presence of dominant species that consistently colonizing islands.

The fitting procedure for the Arrhenius law naturally yields the exponent *z*, which we compare between the homogeneous and heterogeneous cases in Fig. 4. In the region where the Arrhenius model provides the best fit for the homogeneous case, *z* varies between 1 and 0.45, while in the heterogeneous case, *z* varies between 1 and 0.4. The expected value of *z* = 1 arises for very small values of *c*_out_ (see A.F). Moreover, for larger values of *c*_out_, *z* remains higher in the heterogeneous case, and the uncertainty in its estimation is significantly lower than in the homogeneous scenario. This reduction in uncertainty suggests that a heterogeneous species pool stabilizes the species-area relationship, making predictions more robust across different colonization rates.

### 3.3 ISAR in high within-island colonization scenario

We now consider a scenario where within-island colonization exceeds local extinction *c*_in_ *> e*, ensuring that species can establish and persist on islands without continuous immigration from the mainland. This scenario is particularly relevant for islands that are ecologically similar to the mainland or have high connectivity, where species interactions facilitate long-term persistence. To explore this case, we set *c*_in_ = 1 and *e* = 0.8, meaning that local recolonization processes dominate over extinction events. As in the previous section, we assume a homogeneous species distribution on the mainland.

To determine the ISAR, we compute species richness at equilibrium, *S*_∞_, as a function of island size *n*, using the equilibrium condition for *p*_0_ (Eq. (5)) and the species richness equation (Eq. (6)). Unlike the case where *c*_in_ *< e*, the approximation (8) is no longer valid. Instead, we must compute species richness directly from the equilibrium solution. The only limitation is that numerical computations become challenging for large values of *n*, restricting our exploration to moderate island sizes (*n <* 150). The resulting ISAR curves are shown in Fig. 5a. We observe that the saturating effect also occurs for *c*_in_ *> e*, however, an important difference emerges at low values of *c*_out_: the ISAR curve becomes convex instead of concave. This has a significant impact on the estimation of the Arrhenius exponent *z*, which can exceed 1 in this regime (Fig. 5c), contrasting with the typical values observed when *c*_in_ *< e*. A higher *z* exponent suggests that species richness increases more steeply with island size. This deviation from the expected power-law behavior implies that within-island colonization plays a crucial role in shaping the ISAR, reinforcing the idea that species persistence is not solely driven by mainland immigration but also by local dynamics.

**Figure 5:**
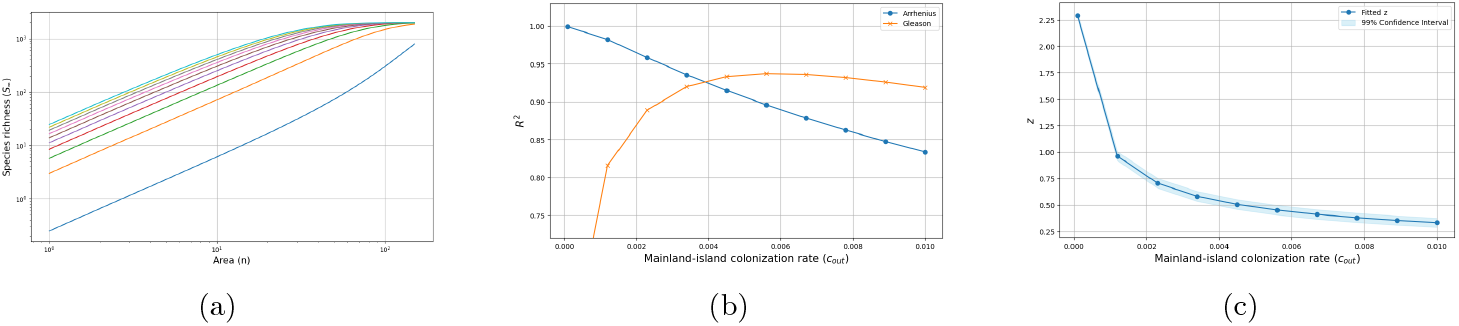
Prediction of species-area relationships with the multiscale metapopulation model (3) under local dynamics with *c*_in_ *> e*. In Fig. (a), the island species-area relationship is shown for different values of the mainland-island colonization rate (*c*_out_), illustrating how species richness (*S*_∞_) varies with island area (*n*). Species abundances (*a*_*m*_) are assumed to be uniform in the mainland. The parameters are *M* = 2000, *c*_in_ = 1, and *e* = 0.8. The mainland-island colonization rate *c*_out_ varies from 10^−4^ (bottom curve) to 10^−2^ (top curve), highlighting how increasing dispersal from the mainland influences species accumulation on islands. In Fig. (b), the coefficient of determination *R*^2^ is shown for the statistical fit of the Arrhenius (1) (blue) and Gleason (2) (orange) laws to the predictions in Fig. (a), as a function of *c*_out_. Each point represents a specific mainland-island colonization rate, ranging from 10^−4^ to 10^−2^, illustrating how well these classical laws approximate the species-area patterns predicted by the multiscale metapopulation model in the homogeneous case. In Fig. (c), the exponent *z* is obtained from the statistical fit of the Arrhenius species-area relationship. For each estimated value of *z*, a 99% confidence interval is plotted, providing an estimate of the statistical uncertainty in the exponent determination.

The behavior of the Arrhenius and Gleason models remains qualitatively similar to the *c*_in_ *< e* case, as shown in Fig. 5b. However, quantitatively, the fit of both models deteriorates, with a lower overall performance in capturing species richness patterns.

## 4 Discussion

We refer to metapopulation theory to define networks of habitat patches within an island, where each patch can host at most one population of a given species. Metapopulation dynamics have been extensively studied during the last decades [15, 17], and the relative colonization and extinction rates across habitat patches should determine (i) whether the species can survive or not in the metapopulation, and (ii) expected population density across patches, conditional on persistence. Here, we further consider that colonization from an external mainland can happen, introducing two distinct colonization parameters: one for within-island dynamics (*c*_in_), and another for immigration from the mainland (*c*_out_). We find that when *c*_out_ is small enough, like in many contexts of oceanic islands, the expected number of species with at least one population (i.e., island richness) varies with island size, in terms of the number of habitat patches in the island, and follows the classical power-law relationship. This represents a dynamical equilibrium [39], as species in the local metapopulation can go extinct while new immigrants may arrive and establish from the mainland. The emergence of the Arrhenius law in a context of multiscale metapopulation dynamics is a new result per se, offering valuable insights into the mechanisms driving spatial biodiversity dynamics.

The connection between Island Biogeography and Metapopulation theory is natural, as both consider the roles of colonization and extinction dynamics in species presence or absence [34, 16]. Ovaskainen & Hanski [34] used the Incidence Function model to characterize the local probability of species presence, depending on connectivity and the occupancy of potential sources, assuming independent species dynamics and that islands can include a single population of each species. We advocate that large islands can host multiple populations, promoting a multiscale metapopulation perspective. Our model is the first to combine finite metapopulation dynamics within islands and the contribution of colonization from an (infinite) mainland [20, 1], in a multispecies context. Our work sheds new light on metapopulation dynamics by considering habitat structure and colonization dynamics at multiple spatial scales, extending previous analyses [20, 1]. While traditional metapopulation models focus on the dynamics and occupancy of a single species, we introduce a simple multispecies extension, assuming independent colonization-extinction dynamics for each species. In this sense, our approach aligns with neutral metacommunity theory, akin to the mainland-island model of Hubbell [19]. Metacommunity theory, based on island biogeography, emphasizes the effect of community size on diversity patterns. The key distinction between the neutral metacommunity framework and a multispecies metapopulation model lies in birth-death dynamics: in the metapopulation model, populations appear and disappear irrespective of individual dynamics, whereas metacommunity theory considers individual birth-death dynamics within local communities. The applicability of these models depends on the realism of their assumptions. The popularity of the metapopulation model in the 90s sometimes led to misleading applications, such as in conservation. Neglecting internal population dynamics can be unrealistic in many cases [4, 11], but the metapopulation approach remains useful for spatial ecology and conservation in many contexts. Our results emphasize that colonization-extinction dynamics at multiple spatial scales have been underexplored, yet they can help predict basic patterns like the SAR/Arrhenius law. While the independence of species dynamics in our model may seem unrealistic, it may hold in small and remote islands where habitat carrying capacity is not the primary limitation. Furthermore, island biogeography theory suggests that negative interactions are weaker, which can influence plant and animal ranges [24] and predator defenses [38]. However, this assumption restricts the model’s applicability to island conditions that meet these criteria.

One of the main concurrent of the Arrhenius law has been the semi-log relationship proposed by Gleason [13]. A major argument of Gleason against Arrhenius law was the fact that the power law increases to infinity with island area, which is unrealistic. However, the range of island sizes is usually limited and in practice most of the data indicate a monotonous increase of richness with island size, without reaching an asymptote [3]. Our results bring some novel insights on the issue. If the colonization rate from the mainland to island is large enough, we find that we can reach a state where all species from the mainland can establish and persist over a long term in the islands, leading to an asymptote in ISAR equal to mainland richness. In addition, we show that the power law in general provides a better fit than the semi-log relationship when the colonization from the mainland is smaller, while the semi-log relationship can provide a better fit at larger *c*_out_ values (Fig. 5). In addition, the values of the exponent of the power law found with our model include the range of values, aka, *z* = 0.2 − 0.4, generally found in real ISAR [12]. The range continues to extend upward, reaching *z* = 1 as *c*_out_ approaches zero, and even beyond when *c*_in_ *> e*. In our case, realistic values of *z* stand for large enough *c*_out_ values, suggesting that our model can be supported by empirical evidence only in this case. However, *c*_out_ must still be not too large enough to avoid richness reaching an asymptote and deviating from the power law (Fig. 5). The range of *z* values should also depend on the shape of species abundance distribution in the mainland, as shown in the comparison of homogeneous abundances and log-series distribution in the mainland (Fig. 4). Further research will be needed to address the influence of other Species Abundance Distributions in the mainland.

Our approach is in line with the classical biogeographical framework that states that the immigration-extinction balance shapes the relationship between richness and area without considering any other factor. Much discussion in the literature has focused on the simplistic nature of the model and the processes it can acknowledge [25]. There are two main kinds of limitation. First, the original Island Biogeography Theory neglects the influence of ecological niche differences among species, and how the physical environment in the islands can in fact select only species adapted to the available ecological conditions. Then, the diversity of environmental conditions in the island can matter more for island richness than its geographical extent. In fact, many studies have pointed out that habitat diversity can be a better prediction of richness than geographical area [36]. In the context of our metapopulation model, we used the number of habitat patches in an island as a measure of its “area”. It is then important to note that our measure of island size is not necessarily a proxy of geographical area. We can consider that a given island includes different kinds of habitats and use our metapopulation model to represent spatial occupancy in each habitat separately. Yet it remains clear that our model is “neutral” in the sense that it does not acknowledge the influence of species adaptation to local environmental on its establishment and persistence. In addition, it does not consider the role of interactions between resident species. Recently, several lines of research considered the possible roles of niche-based processes.[35] suggested that understanding biodiversity patterns in a multiscale perspective needs integrating local ecological interactions with large-scale dispersal dynamics. Competition across species can play a role, and when the carrying capacity of a habitat is reached, it can prevent establishment of new species in this habitat [42]. Finally, the Trophic Island Biogeography Theory has been proposed to acknowledge the role of trophic interactions on biodiversity patterns in islands [14, 29]. For instance, the establishment and survival of a predator is only possible if preys are present in the island. Then the immigration probability of the predators depends on the prior presence of preys, and its extinction probability also depends on the fact that preys persist or not in the island. The structure of the trophic network can then greatly influence the richness of species at different trophic level, and alter the predictions of the initial MacArthur and Wilson’s model [27].

A promising extension of our work would be to integrate species interactions, such as competition or trophic dependencies, into the model. Recent advances in trophic island biogeography [38] suggest that predator-prey interactions can significantly influence island species richness patterns. Competition can also limit colonization success, particularly when habitat carrying capacity is limited [40, 42]. These dynamics have been explored in competition-colonization trade-off models and Lotka-Volterra interaction frameworks, which could be integrated into future extensions of our model [30]. Additionally, positive interactions, such as mutualistic dependencies, could further shape SAR patterns, as suggested by recent work on facilitation-driven metacommunities [32]. Future research could also explore more complex spatial structures, such as island-to-island dispersal networks [20].

In conclusion, our study introduces a multiscale framework for understanding biodiversity dynamics in fragmented landscapes. By integrating metapopulation theory and island biogeography, we provide fresh insights into island species-area relationships. While our model simplifies species interactions and habitat diversity, it offers a useful tool for predicting species richness in isolated systems and lays the foundation for future extensions that could incorporate more complex ecological dynamics.

## Author contributions

F.M. initially conceived the framework. M.C., F.M., and E.P. wrote the first draft of the manuscript, developed the theoretical model, and carried out the analyses and interpretation of the results. M.C. performed the numerical simulations. All authors contributed to the conception of the study and to the revision of the manuscript.

## Acknowledgements

We gratefully acknowledge funding from GDR TheoMoDiv for supporting the organization of a workshop.

## Script and code availability

Script, codes and figures are available online on Github [6]. The code is written in Python.

## A Simulation details

To analyze species-area relationships (SAR), we compare two classical models: the Arrhenius and Gleason models. The Arrhenius model describes species richness as

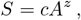

where *c* and *z* are parameters that determine the scaling relationship. In contrast, the Gleason model follows the form

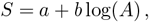

where *a* and *b* are parameters to be estimated. These models offer different perspectives on how species richness scales with area, and fitting them to simulated data allows us to evaluate their applicability under different ecological scenarios.

Our objective is to fit species richness data to these models using data generated from the multiscale metapopulation framework. When the conditions *c*_in_ *< e* and *β* ≪ 1 hold, we use the approximation in Eq. (8) to generate the data. This approximation provides a way to compute species richness *S*_∞_ as a function of island area *n*, allowing us to systematically generate data points for different island sizes *n* and varying distances from the mainland *c*_out_.

In scenarios where *c*_in_ *> e*, the approximation (8) is no longer valid. Instead, we directly compute the equilibrium solution for *p*_0_ using Eq. (5) and apply the richness equation (6) to generate data. This ensures that the model fitting process remains robust across different ecological conditions.

To fit the models to the generated data, we employed the `curve fit` function from the `scipy.optimize` module in Python. This function uses the Levenberg-Marquardt algorithm, a non-linear least squares method, to adjust the model parameters by minimizing the sum of squared differences between the observed data and the model’s predictions. The process begins with an initial guess for the parameters, which guides the algorithm towards an optimal solution.

The `curve fit` function provides not only the optimal parameter values but also a covariance matrix. The diagonal elements of this matrix represent the variance of the parameter estimates, which are used to compute confidence intervals for each parameter. These intervals quantify the uncertainty in the parameter estimates, providing insights into the reliability of the fitted model.

By fitting the Arrhenius and Gleason models to our data using `curve fit`, we gain insights into the relationship between species richness and area. The confidence intervals provide a measure of uncertainty, enhancing our understanding of the model’s applicability. This approach allows us to assess the effectiveness of each model in describing species richness patterns under various ecological conditions.

## B The deterministic mainland-island metapopulation model

Introduced by Levins in the late 1960s [22], the metapopulation model describes the dynamics of populations in a patchy environment, governed by two key processes: colonization, the capacity of a population to disperse to and establish in empty patches, and extinction, the loss of populations within individual patches. The dynamics of the proportion of occupied patches *p*(*t*) at time *t* is given by:

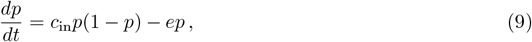

where *c*_in_ represents the colonization rate, and *e* represents the extinction rate.

Building on Levins’ model, an extension was introduced to incorporate the arrival of immigrant species from the mainland, resulting in a mainland-island metapopulation structure [18, 15]. This structure is characterized by a mainland population that acts as a constant source of colonists for an island divided into an infinite number of habitat patches (see Fig. 1a). The deterministic dynamics of the mainland-island system can be described by the following equation:

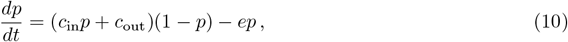

where *c*_out_ is the colonization rate from the mainland to empty patches in the island.

In the limit of a low colonization rate within the island (*c*_in_ → 0), Eq. (10) converges to an equilibrium state given by:

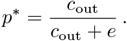

This suggests that some populations will always be sustained through immigration from the mainland, even when the extinction rate is high, so that species extinction in the metapopulation never occurs. This contrasts with Levins’ metapopulation model (9), where population dynamics are influenced by the threshold *c*_in_ *> e*. In Levins’ model, extinction occurs when the colonization rate is insufficient to counterbalance the extinction rate. The mainland-island model (10) can be solved analytically, offering valuable insights into the system’s behavior under different conditions.

The deterministic models (9) and (10) are based on several simplifying assumptions. They assume an infinite number of patches, which limits their applicability to real-world scenarios involving a finite number of habitat patches. All patches are considered equal in size and quality, disregarding variations in habitat suitability or resource availability. Additionally, the models assume that all patches are fully connected by colonization, ignoring the potential impact of distances between patches or isolated patches on system dynamics.

A significant consequence of assuming an infinite number of patches is the absence of stochastic extinctions. In finite systems, random fluctuations in population sizes can lead to extinction events [9], even if the average colonization and extinction rates allow species persistence in the deterministic model. In particular, a key distinction between the stochastic and deterministic models (9) is that the stochastic model predicts global extinction, regardless of the parameter values. By neglecting these random fluctuations, the deterministic model neglects the risk of extinction in real-world metapopulations. In this paper, we analyze a multiscale metapopulation model that explicitly accounts for these random fluctuations.

## C Multiscale metapopulation model for a single species

The multiscale metapopulation model offers a robust framework for investigating the dynamics of populations distributed across multiple patches. Unlike deterministic models, which focus on average population sizes and predict outcomes based on fixed parameters, stochastic models explicitly incorporate random fluctuations in population numbers. These fluctuations arise from various factors, including birth and death events, immigration from the mainland, and local extinctions, introducing uncertainty into the population dynamics. By accounting for this randomness, stochastic models provide a more realistic representation of population dynamics in fragmented or variable environments, where outcomes are shaped by both chance and deterministic processes.

A common approach for modeling stochastic metapopulations is through a continuous-time Markov chain (CTMC). In this framework, the system’s state is defined by the number of occupied patches, with transitions between states occurring randomly over time. The rates of these transitions are determined by the underlying biological processes, such as colonization, extinction, and immigration, which govern the changes in population dynamics across the patches.

The transition rates between states depend on two key processes, represented by *λ*_*k*_ and *µ*_*k*_, for all *k* ∈ [0, *n*], where *n* is the total number of habitat patches in a finite island. First, the rate at which new species occupy empty patches on the island is given by

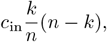

where *c*_in_ is the colonization rate within the island, allowing dispersal from occupied patches to empty patches. Second, the immigration rate from the mainland to the island is given by

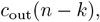

where *c*_out_ is the mainland-island colonization rate, reflecting the distance between the mainland and the island. Mainland colonization is possible only on empty patches, and a higher *c*_out_ increases the
likelihood of colonization events. The parameter *e* represents the extinction rate on the island and is assumed to be independent of the island size *n*. This rate governs the probability that occupied patches become empty due to local extinctions.

The dynamics of the number of occupied patches *k* on the island of size *n* are described by the following transition rates:

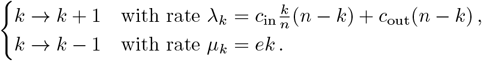

The dynamics of a CTMC are typically described by a set of master equations. The master equations are a set of differential equations that describe how the probabilities of the system being in any particular state change over time. In other words, they govern the dynamics of the probability distribution, which tells us how likely the system is to be in any of its possible states (such as having a certain number of occupied patches on the island) at any given time. Consider an island of size *n*, where *p*_*k*_ represents the probability that exactly *k* patches are occupied in the island. The dynamics of the probability distribution *p*_*k*_ is governed by the set of master equations:

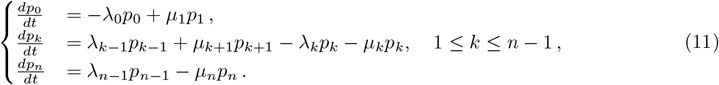

The first equation ensures that the probability of having zero occupied patches, *p*_0_, cannot become negative. The last equation ensures that the probability of having all *n* patches occupied, *p*_*n*_, cannot exceed the maximum capacity, thus enforcing the boundary condition for the system.

Alonso and McKane [1] provided a detailed mathematical analysis of the model (11). They derived the stationary distribution of *p*_*k*_ for *k* ∈ [*n*] and proposed an approximation for the probability of global extinction, *p*_0_, of the metapopulation. This approach offers valuable insights into metapopulation dynamics by integrating regional-scale mainland-island colonization processes with local-scale dynamics of colonization and extinction within individual islands.

Building on this model, Huth *et al*. [20] further incorporated variation in the finite sizes of islands. This extension introduced the concept of rescue effects, where larger or more connected islands could act as refuges, helping populations avoid extinction on smaller or more isolated islands.

## D Limiting cases

### D.1 Isolated island

Consider the limiting case where *c*_out_ approaches zero, leading to a small *β*, effectively isolating the island from the mainland. This scenario is particularly relevant for studying the intrinsic dynamics of the metapopulation with minimal external colonization influence. In this limit, we can show that the probability of extinction, *p*_0_(∞), increases proportionally to 1*/β*. To establish this, we derive an approximate form of Eq. (5) for small *β*.

The Gamma function Γ(*k* + *β*) can be approximated using a first-order expansion around *β* = 0:

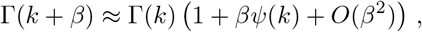

where *ψ*(*k*) is the digamma function, the derivative of the logarithm of the Gamma function:

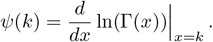

Additionally:

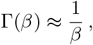

thus, the ratio 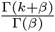 simplifies to:

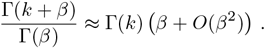

Applying this to the summation in Eq. (5), we obtain:

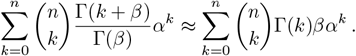

Since Γ(*k*) = (*k* − 1)! for integer *k*, the sum simplifies to:

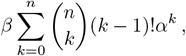

for very small *β*, the probability of extinction *p*_0_(∞) can thus be approximated as:

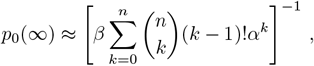

which explicitly shows that *p*_0_(∞) increases proportionally to 1*/β*.

### D.2 No within-island colonization

In the limiting case where there is no within-island colonization (*c*_in_ = 0) but a non-zero mainland-island colonization rate (*c*_out_ ≠0), the equilibrium distribution of the metapopulation model simplifies and converges to a binomial distribution. This result aligns with the findings of [28].

### D.3 Absence of mainland-island colonization rate

The model with no colonization from the mainland (*c*_out_ = 0) and its quasi-stationary state is often referred to as the logistic stochastic model due to its similarities with the deterministic logistic growth model. When *c*_out_ = 0, it is evident from (4) that *p*_*k*_(∞) = 0 for all *k* 0, while *p*_0_(∞) = 1. This result implies that, in the absence of mainland colonization, the probability of metapopulation extinction is almost certainly 1. Even if the within-island colonization rate (*c*_in_) exceeds the extinction rate (*e*), the metapopulation will ultimately face extinction. However, before extinction occurs, it reaches a nonzero quasi-stationary occupancy state, with the time to extinction depending on the colonization-extinction parameters and the metapopulation size [9]. The properties of this quasi-stationary behavior have been extensively studied in the literature [33], but for the specific model considered in this paper, they have only been analyzed by Alonso & McKane [1].

As highlighted by Alonso and McKane [1], the extinction time is defined as the first instance when the population collapses completely. Consequently, even for small values of *c*_out_, there exists a non-zero probability of extinction. As *c*_out_ approaches zero, this extinction probability asymptotically increases to 1, underscoring the critical role of mainland colonization in sustaining the metapopulation.

## E Approximation of the probability of extinction

For a single species, Alonso and McKane [1] demonstrated that an analytical solution for *p*_*k*_(*t*) can be derived under the assumptions of small *c*_out_ and small *k*. When *k* ≪ *n*, the problem simplifies by assuming the following form for the transition rates:

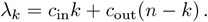

Under these assumptions, the growth rate *λ*_*k*_ becomes linear in *k*, allowing the master equation to be solved analytically using the method of characteristics in combination with the generating function:

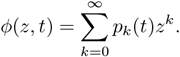

Detailed calculations yield the explicit solution for *p*_*k*_(*t*), with particular interest in *p*_0_(*t*) = *ϕ*(*z* = 0, *t*), given by the exact solution:

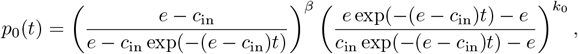

where the initial condition is specified as 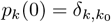, with 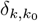 being the Kronecker delta.

## F Approximation of the SAR

For *c*_in_ *< e* and *β* ≪ 1, implying a small *c*_out_, the asymptotic richness given by Eq. (6) can be approximated as a linear function of the island size n:

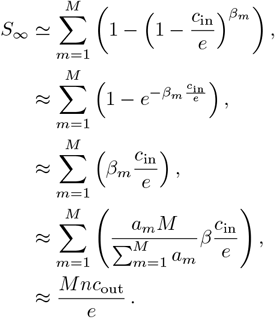

This result follows an Arrhenius-type law with an exponent *z* = 1.

